# BirdFlow: Learning Seasonal Bird Movements from eBird Data

**DOI:** 10.1101/2022.04.12.488057

**Authors:** Miguel Fuentes, Benjamin M. Van Doren, Daniel Fink, Daniel Sheldon

**Affiliations:** Manning College of Information and Computer Sciences, University of Massachusetts Amherst, 140 Governors Drive, Amherst, MA 01003, USA; Cornell Lab of Ornithology, Cornell University, Ithaca, NY 14850, USA

**Keywords:** bird migration, movement ecology, graphical models, big data, species distributions, forecasting

## Abstract

Large-scale monitoring of seasonal animal movement is integral to science, conservation, and outreach. However, gathering representative movement data across entire species ranges is frequently intractable. Citizen science databases collect millions of animal observations throughout the year, but it is challenging to infer individual movement behavior solely from observational data. We present BirdFlow, a probabilistic modeling framework that draws on citizen science data from the eBird database to model the population flows of migratory birds. We apply the model to 11 species of North American birds, using GPS and satellite tracking data to tune and evaluate model performance. We show that BirdFlow models can accurately infer individual seasonal movement behavior directly from eBird relative abundance estimates. Supplementing the model with a sample of tracking data from wild birds improves performance. Researchers can extract a number of behavioral inferences from model results, including migration routes, timing, connectivity, and forecasts. The BirdFlow framework has the potential to advance migration ecology research, boost insights gained from direct tracking studies, and serve a number of applied functions in conservation, disease surveillance, aviation, and public outreach.

## 1 Introduction

The movements of animals span the globe, and movement is integral to behavior, survival, and reproduction. Monitoring movement is particularly important in the face of climate and landscape change, forces that shape how animals interact with their environments (Bauer et al., 2019; Dunn & Møller, 2019). Capturing movement patterns is critical for effective conservation actions, which may hinge on accurate knowledge of animals’ locations and how geographic and environmental interactions change over time (Fraser et al., 2018; Katzner & Arlettaz, 2020). For these reasons, incomplete movement information frequently impedes progress in science and conservation (Fraser et al., 2018; Katzner & Arlettaz, 2020). Often, these challenges arise from constraints on the number of animals that can be monitored, captured, or re-captured in the field; the weight and shape of tracking devices; the number of tracking devices that can be deployed; and the geographic areas that can be adequately covered.

Migratory birds exemplify the challenges facing movement researchers, as well as the urgent need for additional movement information to inform science and conservation. Migratory birds are important indicators of ecosystem health that connect peoples and places in ways few phenomena can. Migrants rely on a predictable series of seasonally and regionally varying resources which, unfortunately, makes them susceptible to rapid global change (Bairlein, 2016; Rosenberg et al., 2019; Sanderson et al., 2006). In North America alone, an estimated three billion birds have been lost in the last half-century, representing nearly a third of the continent’s avifauna (Rosenberg et al., 2019). To conserve migratory birds and study their responses to global change, data and methods are needed that can capture their movements at population scales. For example, a better understanding of the migratory connectivity of different populations of bird species is crucial (Schuster et al., 2019; Webster & Marra, 2005), but detailed connectivity information is lacking for most species. Unfortunately, wireless tracking devices are too heavy for most bird species, limiting the information that scientists can gather on their movements (McKinnon & Love, 2018). Other sources of direct movement data, such as Doppler weather radars, provide no information on species identities or individual behavior (Bauer et al., 2019; Dokter et al., 2018; Van Doren & Horton, 2018).

Citizen and community science projects provide a source of data on animal occurrence and abundance across the globe. In particular, the eBird database (Sullivan et al., 2014) comprises over one billion global bird observations and has been used highly successfully for population distribution modeling (Fink et al., 2020a; Fink et al., 2020b). Although these citizen science projects are collecting increasing volumes of data across a variety of taxa (e.g. iNaturalist, camera trapping projects, etc.), most of these datasets only provide snapshots of occurrence across a population. Without information on the locations of individuals, it is difficult to infer movement from these datasets. Methods that accurately infer movement behavior from large-scale observational data would unlock troves of citizen science data for use by movement researchers and conservation practitioners.

Previous studies have approached modeling movement from observational data by first extensively cleaning the data to correct for variability from the observation process, and then investigating specific quantities of interest like centroid movement or estimated movement speed (Supp et al., 2021). Some recent work has used relative abundance models from eBird Status & Trends to study migratory connectivity. One approach used deterministic models based on the concept of global energy efficiency, in which simulated birds are distributed to optimize both resource acquisition and energy expenditure (Somveille et al., 2021). Another approach used clustering methods along with an assumption of parallel migration to investigate connectivity (Vincent et al., 2022). All these approaches provide valuable but limited lenses to understand migration because they analyze specific aspects of migration and do not model full trajectories that individual birds may take. By modeling full migration trajectories, we are be able to analyze more aspects of bird behavior.

Here, we present BirdFlow, a probabilistic modeling framework that uses relative abundance estimates from eBird to infer movement behavior across the geographic range of a species. Our method builds on previous work on collective graphical models, which reason about individual behavior from aggregate information about a population (Sheldon & Dietterich, 2011; Sheldon et al., 2013; Sun et al., 2015), and on a related modeling framework from private data analysis in human populations (McKenna et al., 2019). More details regarding the connection to this prior work can be found in Appendix A. Inputs to BirdFlow are weekly high-resolution relative abundance estimates produced by the eBird Status & Trends project (Fink et al., 2020a). The output is a trained model that can be interrogated for biological insight, including estimates of migratory paths, timing, connectivity, and forecasting. BirdFlow models can be trained on any species, even those not tracked by eBird, as long as relative abundance models are available. We evaluate the performance of BirdFlow models on several bird species. We show that while performance can be sensitive to hyperparameter settings, these hyperparameters can be set with the use of direct tracking data. We also show that in many cases, hyperparameter settings which work well for one species can be expected to transfer well to other species. Finally, we perform a case study to show how these probabilistic models can produce a range of high-resolution and temporally explicit biological inferences across species’ entire ranges.

## 2 Methods and Materials

Our central goal in developing BirdFlow is to take advantage of the availability of relative abundance estimates produced by the eBird Status & Trends project (Fink et al., 2020a) to model bird movement. The key challenge is that relative abundance information captures the spatial distribution of a bird population as a whole but does not identify individual birds and track them through time. We demonstrate that it is still possible to infer information about the movement of individuals in a population from population-level data. Figure 1 shows the overall process. The key steps are: (1) pre-processing relative abundance estimates to produce weekly population distributions, (2) specifying a loss function that uses weekly distributions along with a proxy for energetic costs to score potential models, (3) selecting a model structure, (4) optimizing the loss function via a numerical procedure to select the optimal model parameters, and (5) validating the trained model.

**Figure 1:**
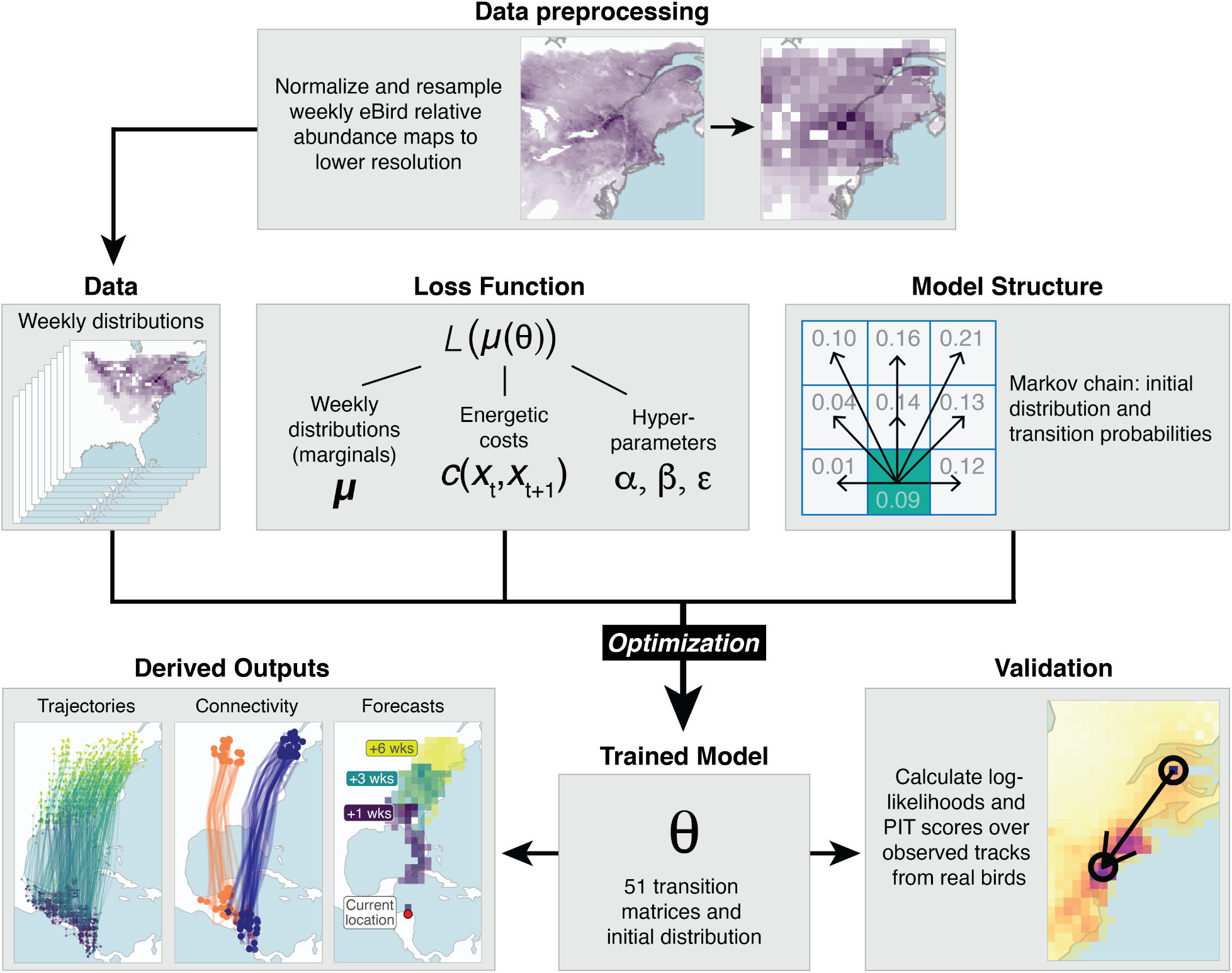
Data preparation and modeling procedure. First, we pre-process eBird Status & Trends data to produce weekly population distributions at a spatial resolution appropriate for the model. We specify a loss function that uses those weekly distributions, along with a proxy for energetic costs, to score potential models. We select a model structure. We fit the model through an optimization procedure to minimize the loss function, producing a trained model. We use observations from tracked birds to evaluate the quality of the model and refine hyperparameters. The final trained model produces various outputs of scientific interest.

Before expanding on those steps, we describe our approach for modeling animal movement as a probability distribution over movement tracks. A movement track is a sequence of locations *x*_1_,…, *x_T_* representing the movement of a single animal over time, recorded at discrete time intervals. Although the methodology supports any time interval, we will use a weekly time interval for the entire paper because that is the time interval of the eBird Status & Trends data. The variable *x_t_* records the location of the animal in week *t* and takes values from a pre-specified finite set 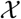 of possible locations, such as the set of all grid cells in a discretized map. For example, if map grid cells are represened by integer identifiers, the movement track 7,14,…, 43 indicates the animal was in grid cell 7 in week 1, grid cell 14 in week 2, and so on, before ending in grid cell 43 in the final week.

We use a probability distribution over tracks to to model the variability in movement behavior among the population of a single species. In this setting, a track becomes a sequence of *random* variables *X*_1_,…, *X_T_* (we use capital letters for random variables) following some joint probability distribution, with the interpretation that each individual in the population follows a movement track sampled independently from this distribution. The probability distribution over tracks defines the *movement model*, which must assign a probability to each possible movement track; for example it could assign probability 0.001 to the track 7, 14, …, 43, which would be written mathematically as Pr (*X*_1_ = 7, *X*_2_ = 14,…, *X_T_* = 43) = 0.001 and means that a randomly chosen bird from the population has probability 0.001 of following this track. There are a total of 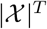 possible tracks, so the movement model can be conceptualized as a vector **p** of 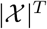 numbers, each of which represents the probability of a different track.

Our representation of a movement model as a distribution over tracks serves as a general-purpose foundation that we will use to link movement distributions to other population-level properties such as the eBird weekly distributions. The model is agnostic to the size of the population, due to the assumption that individuals are independent. One key limitation to this approach is the size of the probability vector: since it grows exponentially in *T*, it will quickly become too large to manage explicitly. For this reason, we must make some assumptions in order to simplify the problem and allow us to efficiently represent **p**. One such assumption, which is commonly used, is to model animal movement as Markovian, so that an animal’s next location is generated based only on the current location, without regard to the past sequence of locations (Patterson et al., 2008). A model with Markovian structure can be encoded by a much smaller vector of probabilities. A Markovian structure is often treated as an assumption about bird behavior. Instead of choosing a model structure based on a model of bird behavior, we construct a loss function which that numerically represents how well a probability distribution models movement, then we determine which model structure is appropriate for that loss function. The processes of constructing a loss function and determining which structure is appropriate for that loss function are explored thoroughly later in this section. We will see that for the loss function we choose, the appropriate structure is the familiar Markovian model.

The movement model is linked to weekly distributions and other population properties through *marginal* distributions. Specifically, the track model **p** specifies all the random variables *X*_1_,…, *X_T_* jointly, from which we can derive the distribution of any subset of those random variables (a distribution of a subset of random variables is called a marginal distribution). So, the weekly distributions we derive from eBird correspond to marginal distributions of the track model. To denote the marginal distribution of a weekly variable *X_t_*, we will use the vector ***μ**_t_*, which has entries that specify the probability that a randomly sampled bird from the population is at a particular location *x_t_* during week *t*. We will index this vector with the following notation ***μ***_*t*_(*x_t_*) = Pr(*X_t_* = *x_t_*). As we will expand on in later sections, an important component of the loss function will be making sure that the track model **p** we learn has marginals ***μ***_*t*_ that match the weekly distributions from the eBird Status & Trends estimates. Note that, in order to simplify the modeling task, we do not consider the uncertainty present in the relative abundance maps or the GPS tracks. The Status & Trends project is clear that the relative abundance estimates are not “ground truth” information about the species distribution, they are estimates with associated uncertainty. Similarly, the GPS tracks have both measurement uncertainty and sampling uncertainty which we do not account for in our validation process. Based on the empirical results, it seems like this omission may be minor.

In the remainder of this section, we will expand on each of the key processing steps illustrated in Figure 1 and outlined above.

### 2.1 eBird Status & Trends Data

We use weekly distributions to train our movement model. Estimating these distributions is a difficult task and a prerequisite to the modeling approach. In the rest of this section, we describe how we use the eBird relative abundance estimates to approximate these distributions. This can be seen in the top row of Figure 1. In the discussion, we describe how the method might be applied to other sources of data and what would be required for compatibility with our method.

The eBird Status & Trends project^1^ estimates the relative abundance of over 600 species at a spatial resolution of 3km x 3km and a weekly temporal resolution (Fink et al., 2020a; Fink et al., 2020b), providing spatial and temporal detail on the seasonally changing population-level abundance patterns of migratory species. We used Status & Trends version 2020, which uses eBird data from 2006–2020 and produces estimates that are broadly representative of that time period. We downloaded relative abundance estimates for 11 bird species that also had available GPS or satellite tracking data (see Table 1 for list of species) as raster files from eBird Status & Trends using the *ebirdst* R package (Auer et al., 2020). Maps of the geographic distributions of these species can be seen in Appendix Figure 10. Among our 11 tracked species, there is substantial variation in the strength of migratory connectivity. For example, Wood Thrush shows “weak-to-moderate” migratory connectivity, and American Woodcocks also show substantial spread in migratory directions while Broad-winged Hawks show relatively strong migratory connectivity (R. A. McCabe et al., 2020; Moore et al., 2021b; Stanley et al., 2021). We chose to use the eBird-based relative abundance estimates instead of the eBird observations directly because the estimates provide a spatiotemporally complete data set by filling spatiotemporal gaps based on modeled relationships with remotely sensed environmental data (Fink et al., 2014; Fink et al., 2013; Johnston et al., 2015) and the estimates remove bias by accounting for systematic patterns of variation inherent in citizenscience observations (Fink et al., 2020b). We loaded rasters at 27 km resolution and re-projected to the Mollweide equal-area projection. Then we aggregated the data to obtain a coarser grid size. This was necessary because of a technological limitation of our compute environment; the GPU that we used to perform the training was limited to grids with about 4000 or fewer cells for a 52-week modeling period. We decided to use as many cells as possible for our experiments, which led to an approximate grid resolution of 100–250 km, depending on the total size of the species’ distribution. The modeling implication of this choice is that weekly movements smaller than the size of a grid cell will not be captured by the model. In the context of migration modeling, this is appropriate because the goal of the model is to capture long-distance trajectories and not local movements. Table 1 shows the average distances covered by the GPS tracks of the 11 species that we model. For each of these species, the GPS tracks cover several times the length of one grid cell, so we are confident that the model can capture migration-level movement. We recommend setting a grid size that captures the relevant movement scale while fitting into the memory of a GPU whenever possible. We used a 110-m resolution shapefile of global coastlines from Natural Earth (naturalearthdata.com) to mask open water, restricting the modeled area to terrestrial environments. Importantly, this restriction allows birds to fly over water, but restricts a bird’s position at the end of a week’s movement to be over land. While this may be restrictive for some species that take extended trips over water, it is not a significant modeling limitation for many bird species. For each week, we normalized relative abundance values by dividing each cell value by the total summed abundance so that the cells sum to one. This gave us “ground truth” estimates 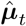 of the weekly distributions, where 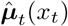 is the fraction of the population that is located in grid cell *x_t_* in week *t* as estimated by eBird Status & Trends (Auer et al., 2020).

**Table 1:** Summary of tracking data used. A “track” is defined as the path of an individual during a calendar year. Individuals monitored for multiple years will therefore have multiple tracks.

| Species | # Individ. | # Tracks | Mean wk/track | Mean km/track | Base ALL | Ref. |
| --- | --- | --- | --- | --- | --- | --- |
| American Woodcock | 67 | 107 | 16.8 | 1334 | -6.52 | <sup>a</sup> |
| Black-bellied Plover | 15 | 38 | 28.3 | 5766 | -5.53 | <sup>b</sup> |
| Broad-winged Hawk | 20 | 35 | 18.1 | 5772 | -5.93 | <sup>c</sup> |
| Blue-winged Teal | 42 | 51 | 21.8 | 2626 | -5.43 | <sup>d</sup> |
| Long-billed Curlew | 91 | 240 | 34.0 | 2163 | -4.86 | <sup>e</sup> |
| Osprey | 230 | 415 | 21.6 | 3501 | -5.74 | <sup>f</sup> |
| Swainson’s Hawk | 43 | 76 | 17.0 | 8443 | -4.96 | <sup>g</sup> |
| Tundra Swan | 50 | 176 | 34.2 | 5325 | -5.42 | <sup>h</sup> |
| Turkey Vulture | 19 | 76 | 35.6 | 4502 | -6.95 | <sup>i</sup> |
| Whimbrel | 32 | 62 | 28.3 | 10884 | -5.66 | <sup>j</sup> |
| Wood Thrush | 20 | 37 | 12.0 | 1859 | -5.93 | <sup>k</sup> |

<sup>a</sup>Moore et al., 2021a, 2021b.
<sup>b</sup>Harrison, 2022.
<sup>c</sup>R. McCabe and Goodrich, 2022; R. A. McCabe et al., 2020.
<sup>d</sup>Ramey et al., 2019.
<sup>e</sup>Carlisle, 2022.
<sup>f</sup>Bierregaard, 2019; Jensen, 2018; Martell and Douglas, 2019; Martell et al., 2001.
<sup>g</sup>Kochert, 1998; Kochert et al., 2011.
<sup>h</sup>Ely et al., 2020.
<sup>i</sup>Bildstein et al., 2014; Dodge et al., 2014.
<sup>j</sup>Tibbitts et al., 2018.
<sup>k</sup>Stanley et al., 2021.

### 2.2 Loss Function

The next step is using the weekly distributions derived from eBird Status & Trends to construct a loss function. A loss function will assign numerical scores to track distributions based on how well they model movement, with the convention that lower scores are better. Later, we will numerically optimize the loss function to find the best possible model under this numerical criterion. One component of the loss will be to ensure that the model **p** has marginals that match the ground truth weekly distributions. This will ensure that the weekly positions of the birds matches what is expected from eBird, but will not alone ensure realistic movements. To ensure that modeled movements are reasonable, the BirdFlow loss function will also include a component that acts as a proxy for the energetic costs associated with moving between locations. To make the notation more concise, we introduce the vectors ***μ*** and 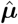, where the vector ***μ*** contains all relevant marginals from the movement model, and the vector 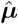 contains all of the weekly ground truth marginals. For example, this allows us to write a function of all the weekly marginals as *f* (***μ***) instead of *f* (***μ***_1_;…, ***μ***_*T*_), and makes the equations more compact without changing their meaning.

We will refer to the loss component that measures the correctness of the weekly distributions as the “location loss”. This loss component is denoted 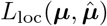 and computes the mismatch between the model’s weekly marginals ***μ*** and the ground truth weekly distributions 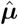 as the mean of the squared error over all of the marginal probabilities:

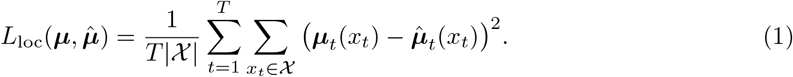

We will refer to the loss component that acts as a proxy for the energetic cost of movement as the “movement loss”. This loss function is denoted *L*_mov_(***μ***) and accumulates the total cost of migration by summing the probability of movement between a pair of grid cells in a given week multiplied by a user-specified cost for that movement, over all pairs of grid cells and all weeks. The probability of moving between a particular pair of grid cells in a particular week is an example of a *pairwise marginal* probability: in the same way that weekly marginals of a movement model encode all information needed to measure the location loss, the pairwise marginals encode all the information needed to measure movement cost. Formally, for any week t, the probability distribution over the pair of locations *X_t_* and *X*_*t*+1_ is a pairwise marginal distribution, and specifies the probability of moving between any pair of locations in that week. We will denote the pairwise marginal distribution by the vector ***μ***_*t,t*+1_, which has entries ***μ***_*t,t*+1_(*x_t_*, *x*_*t*+1_) = Pr(*X_t_* = *x_t_*, *X*_*t*+1_ = *x*_*t*+1_) indexed by a pair of locations *x_t_* and *x*_*t*+1_ that specify the probability that an individual is in location *x_t_* at week *t* and moves to location *x*_*t*+1_ in the next week. With this notation, the equation for this loss function is:

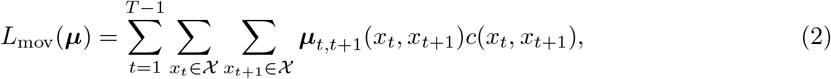

where *c*(*x_t_,x*_*t*+1_) is a user-defined *energy cost* for transitioning from location x_t_ to location *x*_*t*+1_. Note that *L*_mov_(***μ***) is equivalent to the average over the population of the total movement cost incurred by an individual bird during the year. One choice of energy cost is the distance between locations *x_t_* and *x*_*t*+1_, in which case *L*_mov_(***μ***) gives the average total distance moved by an individual. Minimizing this loss function encourages models where birds move the shortest possible distance on average in order to arrive at their migratory destinations. However, we will see later that performance is improved by using a modification with energy cost equal to *c*(*x_t_*, *x*_*t*+1_) = (*d*(*x_t_*, *x*_*t*+1_))^*ϵ*^ a fractional power of the distance *d*(*x_t_*, *x*_*t*+1_) between cells *x_t_* and *x*_*t*+1_, with exponent *ϵ* < 1.0. This energy cost penalizes small distances more than large distances and therefore promotes a model where birds are likely to make fewer large movements instead of many small movements, a behavior that is observed in many bird species (Newton, 2008).

These two loss components form the core of the BirdFlow loss function that is represented in Figure 1. This leads naturally to the optimization problem of searching over all possible track distributions **p** to find one with the lowest score according to this loss function. However, since we wish to optimize over distributions **p** of full tracks but have defined the loss function in terms of the marginals of the track distribution, one more piece of notation is required: we write ***μ***(**p**) to represent the set of marginals ***μ*** derived from the full track distribution **p**; the method for computing the marginals from the track distribution will be described below after specializing to a particular model structure. This leaves us with the following optimization problem:

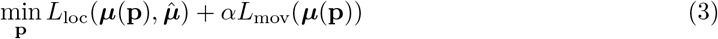

The scalar *α* is a non-negative hyperparameter to control the relative weight of the two loss components.

#### 2.2.1 Entropy Regularization

In preliminary experiments, we found that solving the optimization problem as presented so far leads to a model where birds follow a geographically “narrow” set of optimal paths to exactly minimize movement costs. A real population is expected to have more variability because it will not exactly minimize energy and fitness costs. To address this, we designed an entropy-based regularization term *J*(***μ***), which is added to the loss function to encourage models where birds follow more geographically diverse paths. The details of how this term is computed are given in Appendix B; it is equal to the Shannon entropy of the track distribution (Shannon, 1948), which is a classical measure of the dispersion or uncertainty intrinsic to the distribution. This leads to the following modified optimization problem:

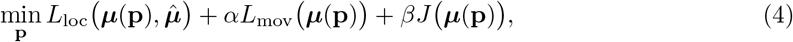

where *β* is another non-negative hyperparameter. Our experiments will show that entropy regularization is important for obtaining biologically realistic movement models, but also that the learned models are very sensitive to the choice of the hyperparameter *β*, with values that are too large leading to track distributions that are so spread out as to be unhelpful. We discuss hyperparameter selection more in the discussion section.

### 2.3 Model Structure

We have now defined a loss function based on the weekly ground truth distributions obtained in the first processing step together with the energy cost and entropy regularization terms defined in the preceding section. Before proceeding with optimization, we must specify the model *structure*, which describes the family of track distributions the optimization procedure will search to find the best model.

In Equation 4, we wrote the optimization problem as searching over all possible track distributions **p**. We will now restrict to searching only over the family of Markov chains, and justify below why this is without loss of generality. A Markov chain is defined by an initial distribution and a sequence of conditional distributions. The initial distribution describes how the birds are distributed in the first week, that is, it specifies Pr(*X*_1_ = *x*_1_) for each location 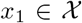. Then, for each week *t* ≥ 1, the conditional distribution Pr(*X*_*t*+1_ = *x*_*t*+1_ | *X_t_* = *x_t_*) specifies the probability that a bird in location x_t_ during week *t* will move to location *x*_*t*+1_ during week *t* +1, for every pair of locations *x_t_* and *x*_*t*+1_. These conditional distributions, which are often called *transition probabilities*, can be used along with the initial distribution to derive all the weekly and pairwise marginals of the model needed for the loss function (Brémaud, 2013; Wainwright & Jordan, 2008).^2^

A key property of a Markov chain structure is that the position of a bird in one week depends only on its position during the prior week, and not any of the earlier weeks. While this may seem restrictive, we will argue that restricting the optimization procedure to search only over Markov chains does not reduce the quality of the estimated model, and is the most principled choice for our problem given the available information. Specifically, note that the loss function only depends on the (weekly and pairwise) marginals ***μ***, and not the whole distribution **p**. Thus, any two track distributions with the same marginals will achieve the same loss, even if they are otherwise different, which makes the optimization problem underdetermined. Conceptually, the optimization can therefore be split into two stages: first, find the optimal *marginals* ***μ***, then use some criterion to select among track distributions **p** with those marginals. A natural strategy is to select the distribution that makes the fewest additional assumptions, the logic being that once we have encoded the relevant information in the loss function, any additional assumptions are unjustified. The *principle of maximum entropy* is a classical embodiment of this strategy; it dictates selecting the distribution with maximum entropy among a set of otherwise equivalent alternatives, which is characterized as being “maximally noncommittal with regard to missing information” (Jaynes, 1957). In our problem, the maximum-entropy distribution with any given set of marginals ***μ*** will always be a Markov chain. This follows from a well known result in the theory of probabilistic graphical models, a class of probability distributions studied in computer science and statistics (Koller & Friedman, 2009), which states that the maximum-entropy distribution with a certain set of marginals is a probabilistic graphical model with a dependence graph in which two variables are connected if and only if they co-occur in one of the specified marginals (Wainwright & Jordan, 2008). In our setup, this means that the maximum entropy distribution will be a Markov chain, since the loss function includes only weekly and pairwise marginals. If we were to use a loss function that depended on different marginals, it would lead to a different model structure for the maximum entropy distribution.

We have now justified restricting our optimization procedure to search only over Markov chains. Let us assume that an arbitrary Markov chain can be represented by a vector ***θ*** of numerical parameters, and, given the parameters ***θ*** describing a particular Markov chain, we can compute the marginals of the corresponding track distribution by a function denoted ***μ***(***θ***). Then, the final form of the optimization problem, now restricted to Markov chains, can be written as:

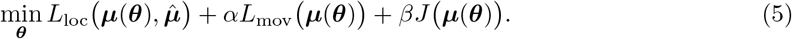

The details of the parameterization of Markov chains, the method for computing its marginals, and the process for solving this optimization problem are described in the next section.

### 2.4 Optimization

We now describe the remaining optimization details, including the Markov chain parameterization and the optimization algorithm. As shown in Figure 1, this process will result in a trained model, which we will proceed to validate.

Let 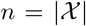 be the number of grid cells. To parameterize a Markov chain, we must specify *n* initial probabilities as well as n^2^ transition probabilities for each of *T* – 1 time steps. To form valid probability distributions, the numbers in each initial and conditional distribution must satisfy the constraints of being non-negative and summing to one. To avoid using an optimization procedure that explicitly enforces these constraints, we re-parameterize each probability distribution using the softmax function *σ*, which transforms an arbitrary vector **u** of *n* real numbers to produce another vector of n numbers that are non-negative and sum to one. Specifically, the ith entry of the output of the softmax function is:

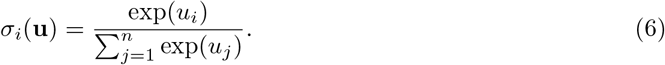

For a matrix *U*, we will also write *σ*(*U*) to indicate the mapping that applies the softmax function separately to each row of *U* to produce a new matrix with rows that are non-negative and sum to one.

We may now parameterize a Markov chain by the parameter vector 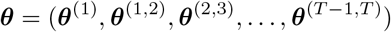, where ***θ***^(1)^ ∈ ℓ^*n*^ is an unconstrained vector of real numbers that determines the initial distribution of *X*_1_, and, for each *t*, the matrix ***θ***^(*t,t*+1)^ ∈ ℓ^*n*×*n*^ is an unconstrained matrix of real numbers that determines the conditional distribution of *X*_*t*+1_ given *X_t_*. The total number of parameters is *N* = *n* + *n*^2^(*T* – 1). We then use the softmax function to transform from unconstrained parameters to probability distributions: the initial parameters ***θ***^(1)^ are mapped to the initial marginal distribution as ***μ***_1_ = *σ*(***θ***^(1)^), and the transition parameters ***θ***^(*t,t*+1)^ for all *t* are mapped to the conditional distributions as 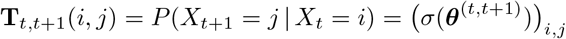. Then, to compute and optimize the loss function, we must also compute the marginals of an arbitrary Markov chain in this parameterization; the procedure ***μ***(***θ***) to compute the marginals from these parameters uses standard Markov chain calculations, and is given in Algorithm 1 in Appendix C.

Because the parameters ***θ*** are unconstrained and the mapping ***μ***(***θ***) from parameters to marginals is differentiable, we can solve the problem in Equation (5) by gradient descent over ***θ*** ∈ ℓ^*N*^. This means that we repeatedly take steps in the direction of the negative gradient of the loss function until we converge to an optimal solution. The model is implemented in Python using the JAX library (Bradbury et al., 2018), which allows us to automatically compute those gradients. We use a gradient descent implementation from the library Optax (Babuschkin et al., 2020) because it is a reliable implementation of a gradient descent algorithm in JAX. Our code, along with a interactive notebook which demonstrates how to use the code are hosted on this GitHub repository https://github.com/Miguel-Fuentes/birdflow. There are other methods to solve Problem (4), for example the proximal algorithm of (McKenna et al., 2019); we selected the gradient descent approach because it is simple and practical. We emphasize that after solving the problem in Equation (5) to obtain the optimal parameters ***θ***, the resulting Markov chain is a global minimizer of the original problem in Equation (4), and has maximum entropy among all minimizers of that problem.

### 2.5 Validation

The quality of a BirdFlow model is sensitive to the choice of hyperparameters, so it is important to validate the model. The most straightforward method of validation is the use of tracking data. We obtained tracking data for the 11 different bird species we fit BirdFlow on from the MoveBank repository (Kays et al., 2022) and other data sources (Table 1). All tracks were obtained with high-precision GPS or satellite tracking devices to ensure minimal uncertainty in location estimates. For Argos data, we retained locations with a location class of 1, 2, or 3, indicating estimated error of <1500 m. For each tracking dataset, we subsampled observations to weekly resolution to match the temporal resolution of eBird relative abundance estimates. To do this, we picked the tracking observation closest in time to the date of relative abundance distribution, as long as the observation was within 4 days of the distribution date. We then matched all tracking observations to the corresponding cell of the distribution raster. When tracking data spanned multiple calendar years, we considered the data from each calendar year as a separate track.

#### 2.5.1 Average Log Likelihood

Once the track data were processed, the primary metric we used to evaluate the model is average log-likelihood (ALL). Given the parameters ***θ*** of a Markov chain in the parameterization described above, let *p*_***θ***_ be the full track distribution described by that Markov chain. Then, for an observed track *x* = (*x*_1_,…,*x_T_*) and parameters ***θ***, the log-likelihood is 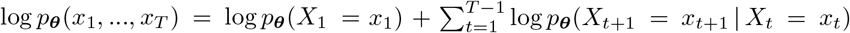. Each of the probabilities in the right-hand side of the preceding equation can be computed easily from the Markov chain parameters by calculations similar to those in Algorithm 1. In practice, many of the tracks span shorter time periods than an entire year and some species have many more tracks than other species. Therefore, to more easily compare results across different species with different numbers of observations, we used the average log-likelihood of bird movements over the total number of observed transitions for that species.

Specifically, each track is split into a collection of movements (*t,x,x*’) where *t* is the starting week, *x* is the bird’s observed location in week *t*, and *x*’ is the bird’s location in week *t* + 1, for each week *t* for which consecutive observations were available. These movements are combined to form the validation dataset 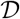. Then, the average log likelihood is given by

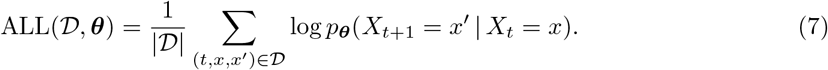

This captures how well the model predicts the movement of the the observed birds and it is comparable for tracks of different lengths and species with different numbers of tracks. Because of this, the average log likelihood is a crucial indicator of model quality. To further contextualize this metric, we constructed a baseline from the eBird relative abundance estimates. The baseline approach ignores the initial position *x* and considers only the log-probability of the destination position *x*’ according to the eBird weekly distribution 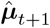

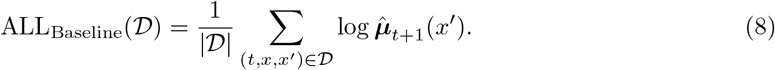

This corresponds to a model where each bird selects a location at random from the weekly population distribution in each week, without regard to its location in the previous week. This “random redistribution” baseline is not biologically realistic, but it captures the information included in the ground truth marginals alone, and can be used to demonstrate how much improvement can be gained by incorporating the information about pairwise marginals through the movement loss function. The values of this baseline for the 11 species we evaluate can be seen in Table 1. Note that an ALL improvement of three nats (the unit for log-likelihood) over this baseline means that the average weekly movement is about 20 times (*e*^3^ ≈ 20) more likely under the model than under the baseline and the average 52-week track is about 1040 times more likely under the model than under the baseline.

#### 2.5.2 Model Calibration

An important capability of BirdFlow is the ability to make probabilistic forecasts. For example, forecasting the distribution of a bird’s location at week *t* + 4 given that it was in a certain location in week *t*. When making forecasts, it is important to understand the model’s *calibration*, or the extent to which the variability of the forecasted distributions matches the observed variability of true outcomes (i.e., a tracked bird’s locations in the future). To measure calibration, we used the *probability integral transform* (PIT) (Gneiting et al., 2007). This transformation uses the cumulative distribution function (CDF) *F* of the forecasted distribution for an eventually observed outcome variable *z*, where *z* is a scalar. If *z* is actually distributed according to the forecasted distribution, then *F*(*z*) will be a uniform random variable; otherwise, the distribution of *F*(*z*) can reveal specific types of miscalibration, such as forecasts being over- or under-dispersed. The distribution of *F*(*z*) is assessed by constructing histograms over many pairs of forecasts and observed values.

We were particularly interested in geographic calibration, that is, the calibration of forecasts of a bird’s location in future weeks given its current location. Since PIT diagnostics apply to scalar quantities, we assessed calibration of forecasts for north-south positions and east-west positions separately. For example, for any grid cell 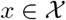, let *u*(*x*) be its east-west position, and let *U_t_* = *u*(*X_t_*) be the random variable for the east-west position of a bird at time *t*. Conditioned on the bird’s location *x* at time *t*, the CDF of the forecast distribution for *U*_*t*+1_ is

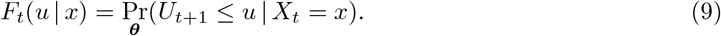

The PIT transform computes the values *F_t_*(*u*_*t*+1_ | *x_t_*) for all observed triples of the form (*t, x_t_*, *u*_*t*+1_) where *t* is a time index, *x_t_* is the bird’s grid cell at time *t*, and *u*_*t*+1_ is the east-west position at time *t* +1.

However, since our map is discrete, we must modify this procedure to correctly account for the probability assigned to discrete outcomes, specifically, the nonzero probability that *U*_*t*_ = *u* in Equation (9). For discrete variables it is common practice to use the *randomized PIT* transform

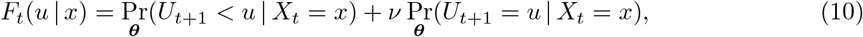

where *ν* is a random variable chosen uniformly in [0,1]. This randomized PIT is evaluated in the same way as the standard PIT.

Since each observed value *F_t_*(*u*_*t*+1_ |*x_t_*) should be uniformly distributed, we can make a histogram of these values and check for uniformity. We followed the same procedure to evaluate north-south calibration, the only difference is that we use the north-south position *V_t_* = *v*(*X_t_*) of the grid cell instead of the east-west position *U_t_* = *u*(*X_t_*).

### 2.6 Experiments

We conducted experiments to assess BirdFlow’s predictive performance, comparisons to baseline models, and sensitivity to hyperparameters.

#### 2.6.1 Hyperparameter Grid Search

We addressed several questions by performing a grid search of model hyperparameters and evaluating the resulting models. The three hyperparameters we are interested in are *α, β* and *ϵ* (the weights on the movement loss *L*_mov_, the entropy regularization term *J*, and the distance exponent used within the energy cost function *c*, respectively). Initial experiments showed that the model is less sensitive to the choice of *α* than other hyperparameters and that a value of *α* ≪ 1 consistently performed well. So, to reduce the search space, we fixed *α* = 0.005 and trained models with different values of *β* and *ϵ*. Conceptually, this places a very high relative weight on the location loss function, which means that BirdFlow weekly distributions will closely match the eBird estimates; then, subject to that “constraint”, the model will minimize the movement costs and entropy costs. We trained the model using every combination of values for *β* ∈ {0.0,0.001,0.002,0.003,0.004,0.005,0.006} and *ϵ* ∈ {0.1, 0.2,0.3, 0.4,0.5, 0.6,0.7, 0.8,0.9,1.0}. We believe this range captures most reasonable values for these hyperparameters because none of the models perform best with the extremal values and the performance appears to vary smoothly as the hyperparameters change. We compared the average log-likelihoods of the resulting models to determine which settings of the hyperparameters led to models that best explain the observed tracks and to understand how hyperparameters affect model quality.

The first question we investigated with the grid search results is the effect of the entropy regularization term and the distance exponent on model quality. We performed an ablation study that compares four model configurations for each species. We compared models with no entropy regularization to models with entropy regularization and models with distance power equal to one to models with distance power less than one. This lets us evaluate how impactful those components are for model quality in isolation and also together.

The second question we investigated with the grid search results was the sensitivity of the model to the choice of hyperparameters. We examined model performance across two methods of hyperparameter selection. First, we tuned each species model by determining hyperparameter values that gave the best average log likelihood for that species; we refer to these as “tuned” model settings. Second, we examined how well each species model performed using hyperparameters chosen based on performance on all *other* species, excluding the focal species. These “leave one out” (LOO) parameters for a species are the hyperparameter values from the grid search results that give the best average log likelihood across all other bird species. We then compared performance using both methods of hyperparameter selection. In particular, wanted to know whether the LOO settings performed well, or if species-specific tuning was required for acceptable performance.

#### 2.6.2 Entropy Calibration

The entropy regularization term is key to the calibration of model predictions. If we increase its weight, the joint marginals will become more diffuse and predicted tracks will have greater directional dispersion. In reality, a given species will show an intermediate level of dispersion. Thus, we can predict that too low settings for the entropy regularization weight parameter will lead to underdispersion and too high settings will lead to over-dispersion. In preliminary experiments we observed that the trend of going from under-dispersion to over-dispersion held for all species, but the value of the inflection point varied by species. As an example, we computed the PIT score for each of the transitions for the American Woodcock (*Scolopax minor*) under several versions of the model and plotted the score in a histogram. A convex-shaped histogram indicates under-dispersion, and a concave-shaped histogram indicates over-dispersion. A uniform (flat) histogram indicates optimal dispersion and a well-calibrated model.

#### 2.6.3 *k*-Week Forecasting

We also investigated model performance for the task of *k*-week ahead forecasting for *k* > 1 to understand how prediction accuracy decreases with time horizon. The procedure for computing the average log-likelihood was slightly modified to compute the average log-likelihood for forecasts *k* weeks into the future. Instead of splitting the tracks into bird movements in consecutive weeks, tracks were split into positions of a single bird *k* weeks apart, that is, we created a data set 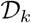 with triples of the form (*t, x, x*’) where *x* was the bird’s position at time *t* and *x*’ was its position at time *t* + *k*. Then, the model and baseline were evaluated on how well they predicted these positions. These modified average log likelihoods were computed as follows

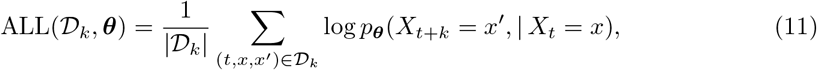

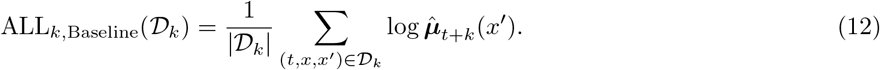

## 3 Results

We now present results of the model validation experiments. Figure 2 shows the results of the ablation study comparing the performance of different model configurations on tracked wild birds. All BirdFlow model types performed better than a baseline model that incorporated only species relative abundance. Models with non-zero entropy regularization and tuned distance penalty exponent (*ϵ*) performed best overall, followed by models with entropy regularization and *ϵ* =1.

**Figure 2:**
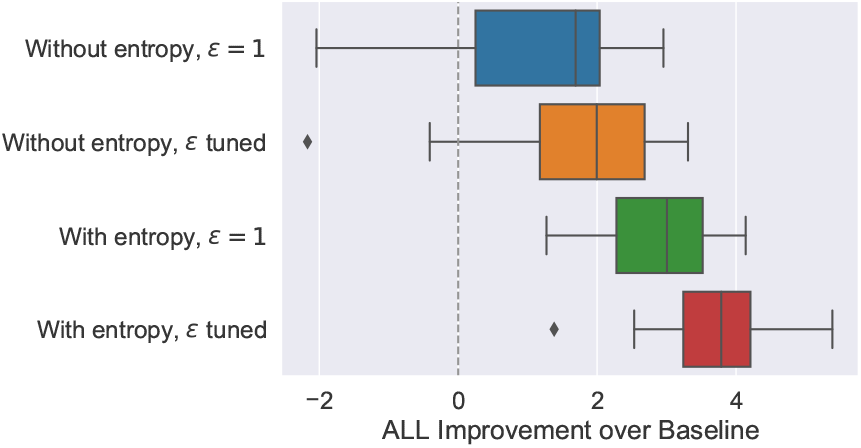
Model type ablation study. For each model version, the distribution of performance (average-log likelihood) improvement over the baseline model for 11 species is displayed as a box and whisker plot. The vertical dashed line marks no improvement over the baseline. Whisker length is at most 1.5 times the interquartile range, with outliers shown as diamonds.

Figure 3 assesses sensitivity to hyperparameters. For most species, the “leave one out” (LOO) parameters, which were selected using only the validation tracks from *other* species, performed nearly as well as models tuned using tracking data from that species. The difference in average log-likelihood between the LOO parameters and the tuned parameters is small compared to the difference between either setting and the baseline. The most notable exception is Swainson’s Hawk, where the LOO parameters perform much worse than the tuned parameters.

**Figure 3:**
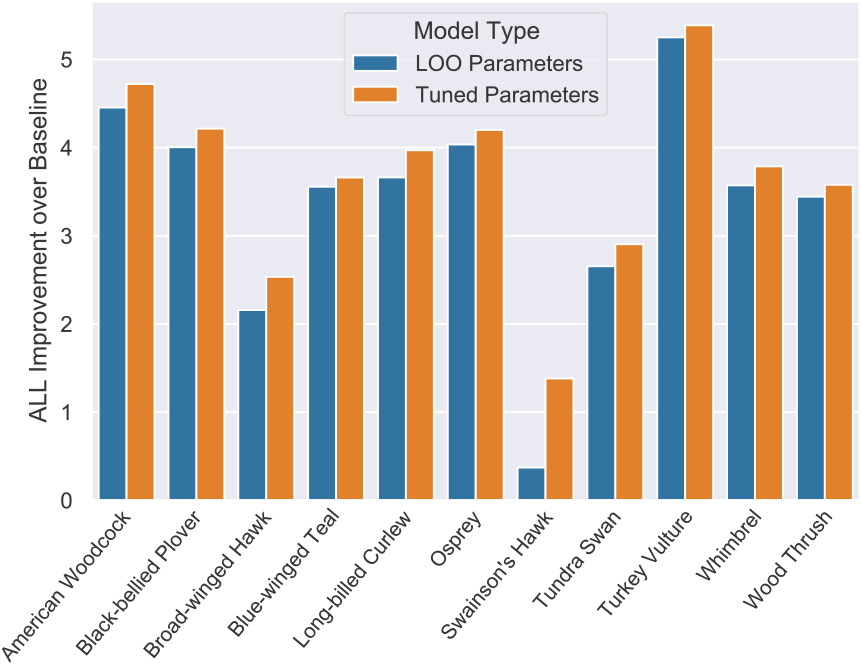
Parameter sensitivity. Bars show improvement over the baseline model for Bird-Flow models with “leave one out” parameters (selected using validation data from other species) vs “tuned” parameters (selected using validation data from the target species). Performance is measured as average log-likelihood of one-week transitions.

Figure 4 shows the effect of entropy regularization on model calibration, which was substantial. PIT histograms for five versions of the American Woodcock model are shown, the distance exponent (*ϵ*) fixed to 0.3 because that is the optimal setting for this species (in terms of log likelihood) and the entropy regularization weights varied since that is the parameter of interest for this experiment. The PIT histograms are closest to uniform for entropy weights of 0.0005 and 0.001, which indicates the best model calibration. Entropy weights that are higher or lower strongly negatively impact calibration. With zero entropy, too many observations occur at the extremes of the forecast distribution, which indicates underdispersed forecasts. With high entropy, too few observations occur at the extremes of the forecast distribution, which indicates overdispersed forecasts.

**Figure 4:**
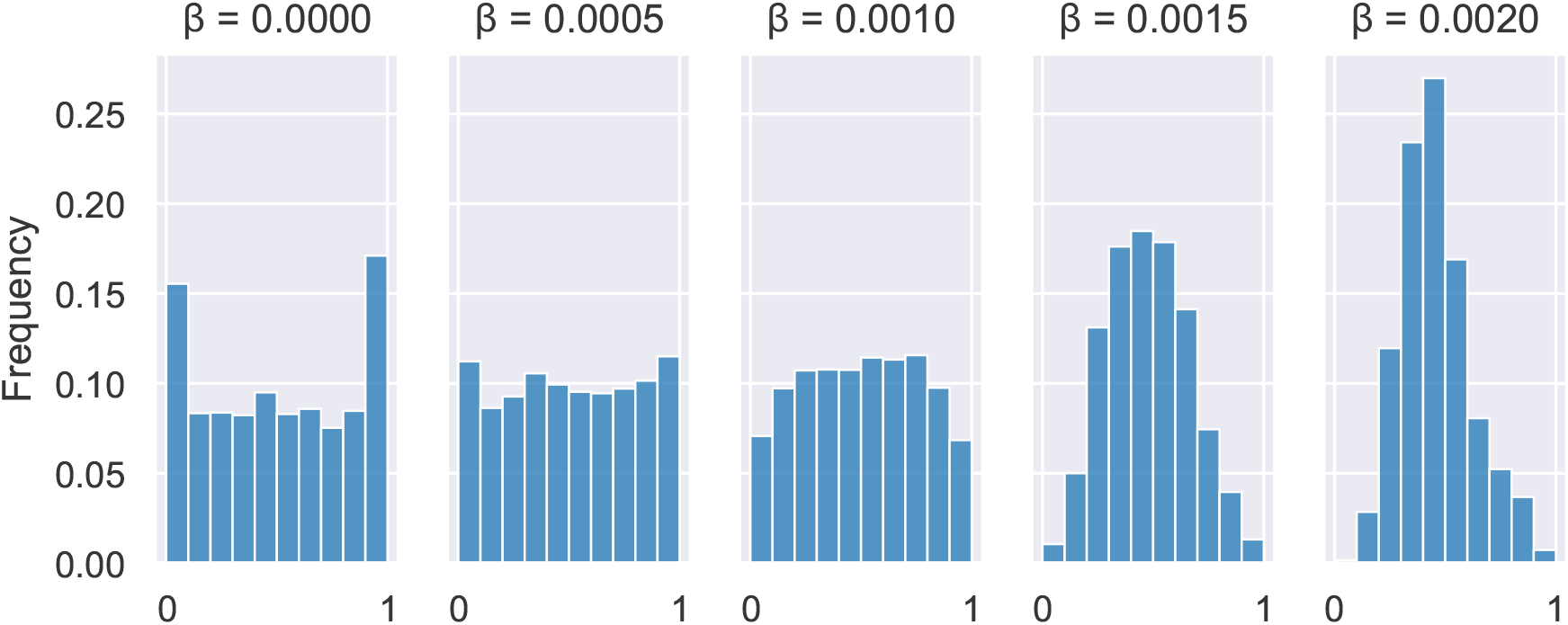
Effect of entropy regularization on model calibration. Randomized PIT histograms are shown for 1-week ahead forecasts of American Woodcock east-west positions for models trained with different entropy weights. Histograms that are nearly uniform indicate well calibrated models.

Figures 5a and 5b show model performance relative to forecast horizon (in weeks). We identi-fied the best-performing model from the hyperparameter grid search (using average log-likelihood) for every species and evaluated the improvement over the baseline for k-week-ahead average loglikelihood for all forecast horizons *k* from 1 to 17. Figure 5a displays those results for each species. For every species, the improvement over the baseline decreases with k. However, there is substantial variation: some species continue to perform substantially better than the baseline up to a forecast horizon of 17 weeks, while others approach the performance of the baseline.

**Figure 5:**
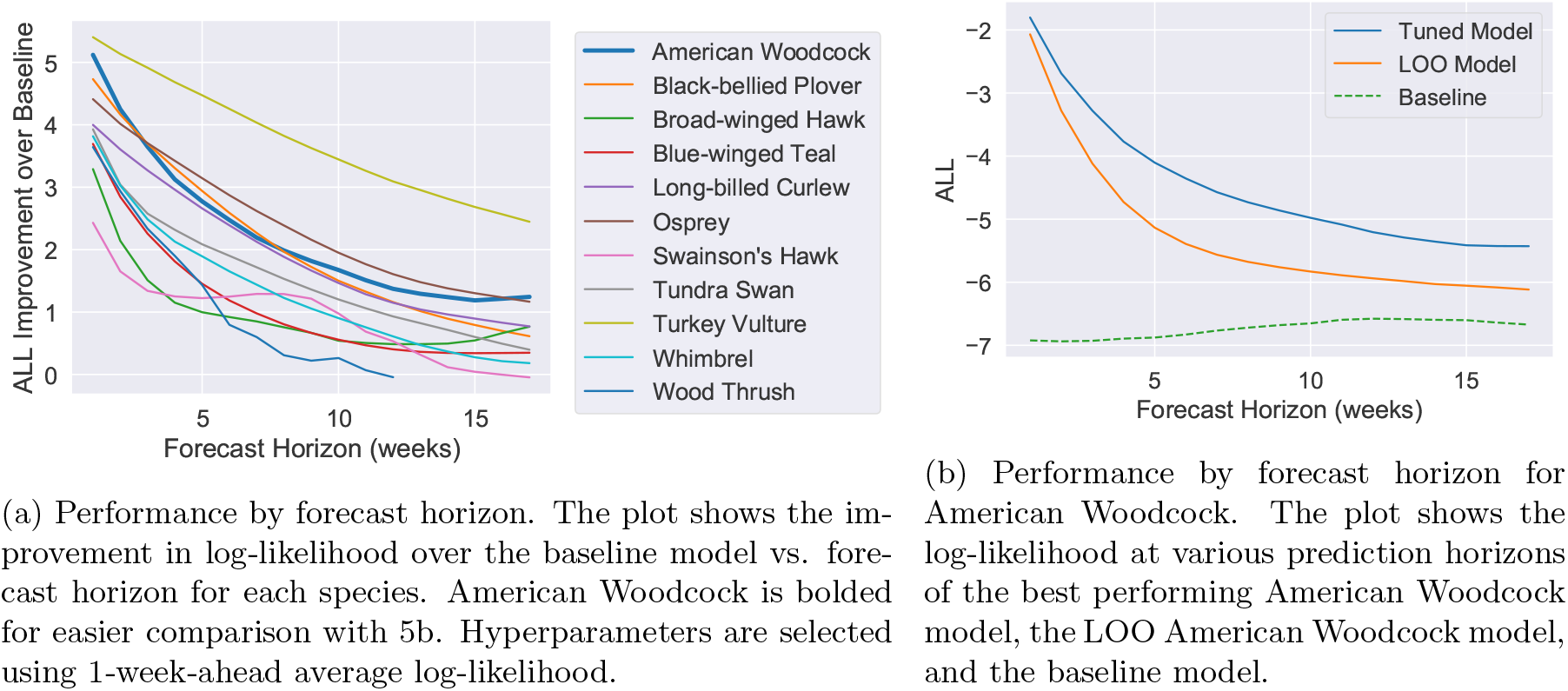
Forecasting performance plots species. We chose this species because we had high-quality validation data from GPS-tracked birds (Table 1). In order to select the hyperparameters, we performed a finer grid search around the best parameters from the original coarser grid search. We selected the model from the finer grid search with the best average log likelihood. We then demonstrate two powerful capabilities of the trained woodcock flow model: sampling and forecasting.

We also compared the tuned woodcock parameters to the LOO woodcock parameters and the baseline in an absolute sense (Figure 5b). The gap between the tuned parameters and the LOO parameters is small at first, but increases with forecast horizon, which indicates that the tuned model performs better relative to the LOO model at larger horizons. Both models performed better than baseline model at all prediction horizons tested.

## 4 Case Study

The experiments investigated various aspects of the model’s performance across a wide range of species. For that investigation, we performed a relatively coarse hyperparameter search. However, we want to demonstrate that if particular care is taken to find hyperparameter settings that work well for a species, the resulting model can output many useful results. To show this, we generated and evaluated model outputs for American Woodcock in addition to what we did for the other

Sampling refers to the generation of synthetic trajectories from the distribution learned from the model. For this demonstration, we simulated 5000 migration trajectories, representing plausible routes of individual woodcocks through the year. The spring portion of a subset of these tracks can be seen in Figure 6a; the color of the line indicates the corresponding time of the year. From these simulated trajectories, we calculated three measures of the spring migration: (1) the distribution of migration departure timing (Figure 6b), (2) the distribution of migration arrival timing (Figure 6c), and (3) the migratory connectivity of breeding populations (Figure 6d). We calculated the distributions of spring migration departure and arrival dates using the *alongTrackDistance* function in the `geosphere R` package (Hijmans, 2017), assessing when each simulated bird moved at least 100 km from its starting location and arrived within 100 km of its ending location. Simulated woodcocks left their wintering grounds between mid-January and early March, arriving largely between early March and early May. To infer migratory connectivity, we used simulated trajectories from the fall migration. We selected trajectories that began in the northwest and northeast sectors of the woodcock breeding range to compare the modeled connectivity of populations originating from different parts of the breeding range. Our model inferred meaningful differences in migratory connectivity between woodcocks breeding in the northeast US and in the Midwest (Figure 6d). The model inferred that woodcocks breeding in the northeast primarily spend the winter in the mid-Atlantic and southeast. In contrast, the model inferred that woodcocks breeding in the Midwest winter primarily along the western Gulf Coast.

**Figure 6:**
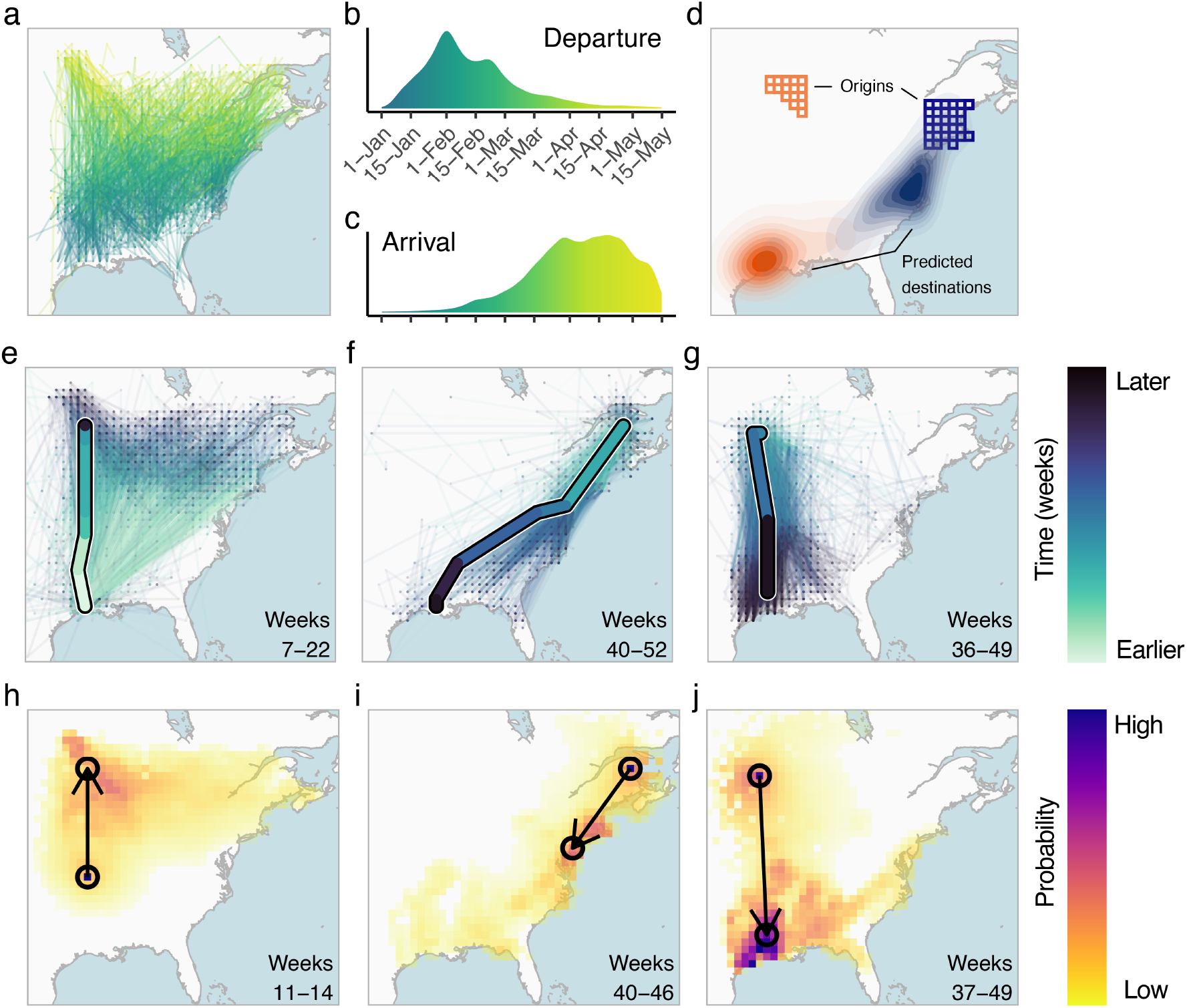
Demonstration of model inferences. Shown are derived model outputs from American Woodcock *Scolopax minor*. (a) Simulated spring migration trajectories (n=1000). (b) Timing of spring migration departure and (c) arrival derived from simulated trajectories. (d) Migratory connectivity: square cells show breeding origins of individuals in the northwest (orange) and northeast (blue) parts of the breeding range. Filled density contours show the predicted wintering distributions of individuals breeding in those respective regions. (e-g) Observed movements of GPS-tracked woodcocks (single thick path) and simulated trajectories (thin paths) for 2500 simulated birds originating at the same starting location as observed birds. (h-j) Conditional forecast distributions: each heatmap shows the predicted movement distribution of a GPS-tracked individual originating within the circle at the base of the arrow. Darker colors indicate a higher predicted likelihood of movement to that area. The point of the arrow shows the observed ending location. Shown are examples of 3-week (h), 6-week (i), and 12-week (j) conditional forecasts.

In addition to sampling tracks from the general population, we can also sample trajectories that start at a particular time and location. To demonstrate this capability, we generated visual representations of modeled tracks alongside actual GPS-tracked individuals to compare modeled trajectories to observed migration routes. For each observed track, we generated 2500 simulated trajectories originating at the same location as the GPS-tracked bird and continuing for the same duration. Then, we plotted observed and simulated routes together in Figures 6e,f,g. The week numbers in the bottom right of these plots give the starting week and ending week of the track which we are simulating. The observed route is given by a thick line while the simulated trajectories are given by thin lines. The observed routes were generally well represented among simulated trajectories. Additional examples of this type of plot are given in Appendix Figure 8.

Forecasting refers to the prediction of a bird’s location in the future based on it’s position at a given time. Our previous experiments analyzed the quality of BirdFlow’s forecasts numerically; for this demonstration we produced visual representations of short-term forecasts. For observed GPS-tracked birds, we took their position at a given start week, then predicted their position at 3, 6, or 12 weeks into the future. Then, we compared the predicted movement forecast to observed movements. This can be seen in Figures 6h,i,j. The numbers in the lower left indicate the start week and end week. The heatmap color indicates the probability the model gives to a particular location. The arrow is directed from the starting position of the GPS-tracked bird to the ending position. The short-term conditional forecast distributions successfully captured observed movements. Additional examples of this type of plot are given in Appendix Figure 9.

## 5 Discussion

Our probabilistic BirdFlow models accurately inferred individual movement behavior using weekly relative abundance estimates from eBird data. For all species studied, our movement model predicted the movements of GPS- and satellite-tracked birds substantially better than a baseline model that included only the weekly species distribution maps. For nearly all species, parameters transferred from other species (LOO parameters) performed well, suggesting that BirdFlow could be used to accurately infer movements without any tracking data inputs in many species. Models fine-tuned with tracking data were most accurate, but the difference between LOO models and tuned models was small compared to the improvements over the baseline model. Overall, the results show that by combining relative abundance estimates derived from eBird data with models of movement costs, it is possible to infer individual movement behavior in a way that is substantially more accurate than baseline models.

### Applications

We show that it is possible to accurately model animal movement from aggregate data. We demonstrate how a trained BirdFlow model can be sampled to investigate migratory routes, timing, and connectivity, and that the model can produce forecasts. In addition to the ecological questions investigated in our case study, samples from BirdFlow models can be used to study other phenomena such as stopover behavior and responses to global change. The experiments show the capability of the BirdFlow model to predict the likely position of a bird several weeks into the future, given a starting time and location. The further into the future the prediction is made, the more uncertainty about the bird’s position accumulates. It is therefore encouraging that the k-week forecasting experiment showed that the model performs consistently better than the baseline even many weeks into the future. The ability to accurately predict the positions of birds could support efforts to monitor the spread of diseases such as avian influenza. Movement researchers with access to even a small amount of tracking data could use our model to infer individual behavior across the species’ entire range—in essence, combining insights from eBird with direct tracks to achieve a more complete understanding of animal movements than either approach can alone. For now, BirdFlow has only been used with eBird data, but the Status & Trends project (Fink et al., 2020a) currently releases abundance estimates for over 1,000 species so movement ecologists can use those data to ask a wide range of questions using a broad range of species with varying movement ecology. Finally, BirdFlow can raise public awareness about biodiversity and ecosystem health by providing a tool for outreach to engage scientists, bird-watchers, policy-makers, and the general public.

### Model Tuning in Practice

Our sensitivity experiment shows that the difference between the LOO parameters and the tuned parameters was usually small compared to the difference between either setting of the parameters and the baseline. However, the results from Swainson’s Hawk indicate that hyperparameter settings will not translate equally well for all species. Further work is needed to fully determine under what conditions hyperparameter settings will transfer well and how to select hyperparameters when no tracks are available. For this reason, we cannot yet provide specific values of the parameters as “default values” which we would expect to perform well for any species. Instead, we advocate for the use of a procedure similar to ours where tracks are used to select models by evaluating key metrics (such as ALL and calibration). Including calibration in the selection process is important because choosing models via their average log-likelihood sometimes favors an entropy weight that produces routes that are more variable than expected. We anticipate that as this method is adopted and tested, the scientific community will develop a set of best practices for validating a BirdFlow model in the presence of some tracks and eventually when no tracks are available. We also believe that the development of these validation practices is a worthwhile endeavor considering the potential applications of these models. These best practices may develop from a better understanding of how the hyperparameters relate to factors such as migratory distance, flight behavior, geographic differences, and others. Of the species evaluated, Swainson’s Hawk migrates the longest distances, with many individual traveling from northern North America to southern South America. We hypothesize that hyperparameters that work well for other ultra-long-distance migrants may transfer better to Swainson’s Hawk.

### Designing Loss Functions

The terms in the loss function that do not depend on data should reflect the biological properties of the target population. We expect loss terms that encode these biological properties more accurately should improve performance. In our case, the movement loss reflects the energy cost of moving and the different values of the distance exponent encodes how much a species will tend to make few large movements compared to many small movements. The entropy regularization term encodes that a real population is not expected to exactly minimize energy and fitness costs, instead showing substantial individual variation in behavior. The ablation study we performed shows that these terms did improve performance, which suggests they are encoding helpful assumptions about bird movement. The addition of an entropy regularization term was crucial for proper model calibration, and using a distance exponent less than one in the movement cost term was important for producing realistic movement patterns. When these components were removed (labeled “Without entropy, *ϵ* = 1” in Figure 2), several species under-performed the baseline. The entropy regularization term seems to be particularly important, because its inclusion alone ensures that the model outperforms the baseline for every single species.

However, this does not mean that these loss components are perfect. Clearly, the energetic cost of a movement depends on more than just distance: it also depends on atmospheric effects and topography. We would expect that constructing an energy cost function that matches the true energetic movement cost more closely would improve performance. As it currently stands, there can be a difficult tradeoff where models trained with low entropy learn distributions that are far too narrow but models trained with a higher entropy learn distributions that send birds in unrealistic directions (see Figure 7). This suggests that there may be a better way to encode biological knowledge about variability in migration paths: intuitively, a high entropy distribution will be very uniform and lead to variability in all directions, but there may be some other loss function that could encourage variability only in desirable directions. Designing loss terms that better encode our biological knowledge is an interesting direction for future work.

**Figure 7:**
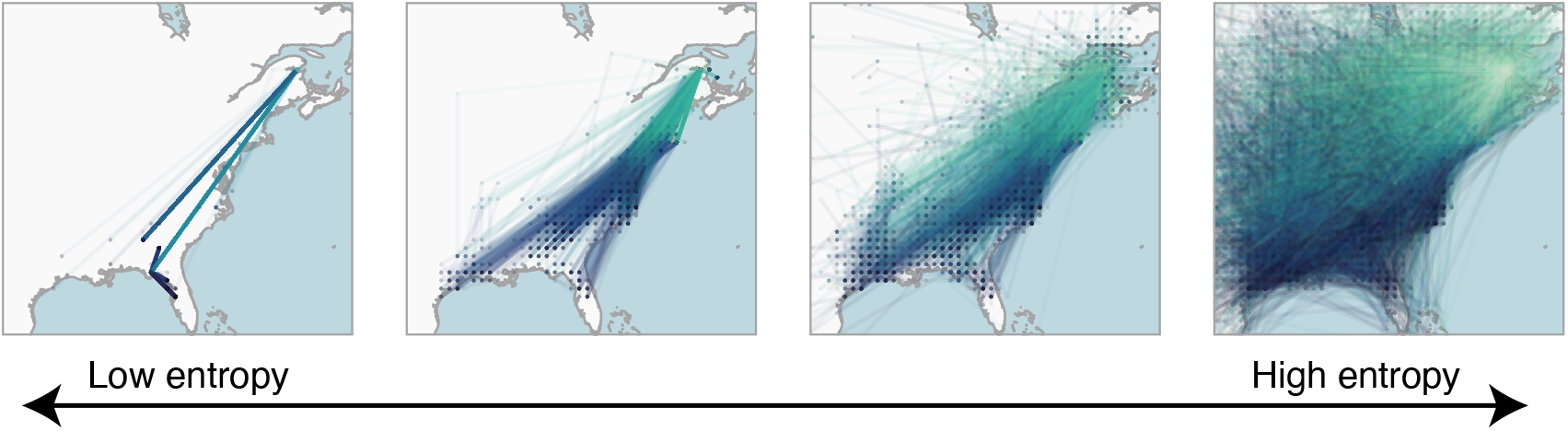
The effect of entropy levels on American Woodcock model samples. The models were trained with the distance exponent (e) fixed to 0.3 and with the entropy weights (0.00, 0.01, 0.02, 0.04). The plot displays 2500 tracks sampled from each model.

### Incorporating other Marginals

In Section 2.3 we explain that our use of a Markov chain to model the track distribution is based on the maximum entropy principle applied to our loss function. While we believe this is a principled choice and we show that it performs well in practice, Markovian models have several known limitations. Because the distribution of future locations depends only on a bird’s current location, the model treats all birds in the same location at the same time identically: their future routes may diverge, but only due to randomness of transitions, and not due to long-term “memory”. This means, for example, that the current implementation of BirdFlow cannot model year-to-year site fidelity. That is, simulated full-year routes are unlikely to return to the same location one year later. For this reason, we currently recommend applying BirdFlow for single migration seasons. For the same reasons, BirdFlow cannot differentiate between individuals of different subpopulations that have different migration strategies but coincide both spatially and temporally. For example, BirdFlow could not correctly model two distinct subpopulations that cross through the same location at the same time. We believe this limitation is minor in practice, because populations with different migration strategies are often separated either spatially or temporally.

These limitations are intrinsic to the Markov chain structure but, if we were to use a loss function that included different kinds of data or encoded different biological assumptions, the choice of model structure would change. For example, we could incorporate site fidelity by adding loss function components that depend on the marginal distribution of a bird’s location at a given time together with its location one year later. If we apply the same logic about maximum entropy to a loss function that also incorporates this “site fidelity marginal”, the resulting model would be more computationally intensive but it would be able to capture site fidelity information in a way that the current model cannot. Constructing loss functions that use other marginals while remaining computationally tractable is another direction for future work.

## 6 Acknowledgements

We are grateful to the eBird Status & Trends team. We thank Tom Auer and Adriaan Dokter for assistance and feedback on our work, and Rob Bierregaard, Autumn-Lynn Harrison, and Michael N. Kochert for permission to use tracking data in this study. This material is based upon work supported by the National Science Foundation under Grant Nos. 1522054 and 1661259. The work of BMVD was supported by a Cornell Presidential Postdoctoral Fellowship. We thank the Leon Levy Foundation; The Wolf Creek Charitable Foundation; NSF DBI-1939187. Computing support was provided by the NSF CNS-1059284 and CCF-1522054, and the Extreme Science and Engineering Discovery Environment (XSEDE) NSF ACI-1548562, through allocation TG-DEB200010 run on Bridges at the Pittsburgh Supercomputing Center. Additional computing efforts were performed with equipment obtained under a grant from the Collaborative R&D Fund managed by the Massachusetts Technology Collaborative.

## A Connection to Prior Work

BirdFlow builds on prior methods for learning a probability distribution from evidence about its marginal distributions. Notably, we previously developed *collective graphical models* (CGMs) (Sheldon & Dietterich, 2011), which are a general formalism for learning the parameters of a probabilistic graphical model from noisy aggregate observations. CGMs were inspired by bird migration modeling (Sheldon et al., 2008), and later used to model human population flows (Akagi et al., 2018; Iwata et al., 2017). Inference and estimation in CGMs is computationally challenging (Sheldon et al., 2013), but many approximations have been proposed (Sheldon et al., 2013; Singh et al., 2020; Sun et al., 2015; Vilnis et al., 2015; Yasunori et al., 2020).

A similar problem setting arises in privacy-preserving data analysis, where noisy aggregate population statistics are released by a central agency such as a census bureau to provide information about population demographics while ensuring privacy of individuals (Dwork et al., 2006). From these noisy, aggregate statistics, an analyst wishes to estimate a full distribution over demographic variables. Private-PGM (McKenna et al., 2019) is a recent algorithmic framework we developed for this setting, which has been successful as a key component of winning entries in privacy competitions (www.nist.gov, 2018, 2020) and of mechanisms for releasing private synthetic data (Cai et al., 2021).

BirdFlow builds on the conceptual underpinnings of Private-PGM, rather than CGMs, to estimate bird movement models. One key difference compared to CGMs is that BirdFlow and Private-PGM ignore sampling variability due to the population being drawn from an underlying distribution. This is appropriate for large populations, where sampling error is smaller in magnitude than measurement noise, and leads to simpler estimation algorithms. A second key difference is that in BirdFlow the model output is a probabilistic model (a Markov chain), while in CGMs the model output is a reconstruction of population flows. While this difference is minor mathematically (one object can be converted to the other), it is a significant practical and conceptual advance to treat the model output as a probabilistic model from which we can construct synthetic routes and create forecasts and many other products. Finally, although CGMs were motivated by bird migration modeling, the current study is the first in-depth examination of the capabilities of any of these methods to accurately model bird migration at this scope, including many species, validation using real tracks, and tuning of of key parameters such as entropy regularization and distance exponent to obtain biologically realistic model outputs.

## B Entropy Regularization

Formally, the entropy regularization term *J*(***μ***) is equal to the negative Shannon entropy of the distribution. The Shannon entropy is a measure of the uncertainty of a distribution, so in order to correct for the model’s overconfidence, we seek to increase the entropy. Since the optimization procedure minimizes the loss function, in order to increase entropy we seek to minimize negative entropy. Hence the loss component being negative entropy. In general, computing entropy requires summing over every possible outcome of a distribution which would be infeasible to do for a distribution of all possible tracks. However, there is a well established result (Wainwright & Jordan, 2008) which states that the entropy of a Markov chain can be written as a function of the entropy of it’s marginals. Using this result, the negative entropy of a Markov chain can be written as

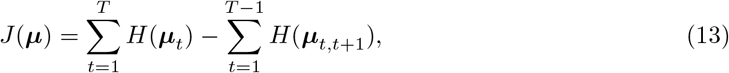

where *H*(***μ***_*t*_) and *H*(***μ***_*t, t*+1_) are Shannon entropies of corresponding marginal distributions, specifically:

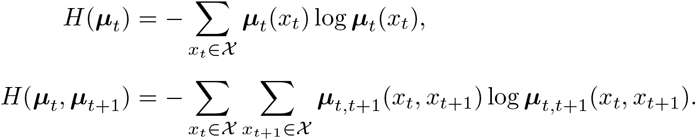

The entropies of these marginals require summing over exponentially fewer possibilities so we can efficiently compute them.

There is a bit of subtlety with our use of this substitution. In the model structure section, we say that the loss function only uses weekly marginals and pairwise marginals and this justifies the use of a Markov chain. But here, we say that the entropy component of the loss function depends only on marginals because we have selected the Markov chain structure. In our view, the location loss and the movement loss constitute the core loss function because they correspond to data and biological knowledge; for these loss components it is always true that they only depend on marginals regardless of the model structure we choose to optimize over. These central components justify the model structure, and once this structure is selected, the entropy regularization is a post hoc addition to correct for overconfidence.

## C Markov Chain Calculation

Algorithm 1 details the process for transforming the parameters ***θ*** into the marginals ***μ***. The parameters ***θ*** are vectors and matrices of real numbers with no constraints, while the marginals and conditional distributions of the model are constrained to be valid probability distributions. Optimizing over a constrained space in inherently more complicated than optimizing over an unconstrained space, so this kind of substitution is relatively commonplace. Importantly, the operations present in this algorithm (exponentiation, multiplication, and addition) are all differentiable. In order to perform optimization via gradient descent, we must be able to compute the gradient of the loss function in terms of the parameters ***θ***. Computing those gradients requires the differentiability of not only the loss function, but also the mapping from parameters to marginals. We omit any manual derivations of gradients because modern automatic differentiation software (like JAX) computes these gradients automatically.

### Algorithm 1: Differentiable mapping from parameters ***θ*** to marginals ***μ***

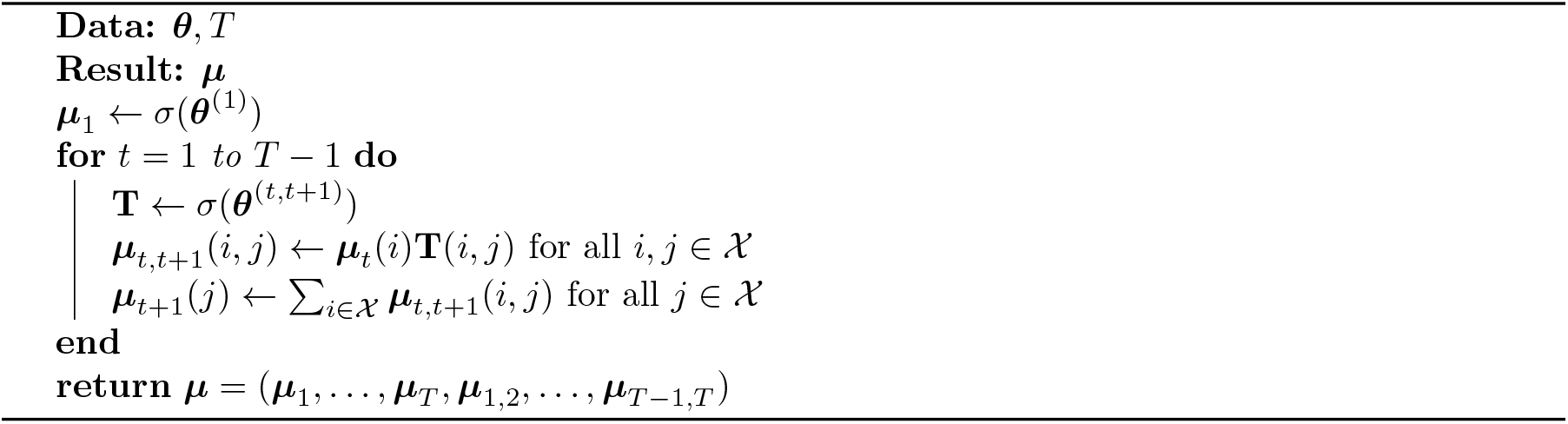

## D Additional Case Study Figures

**Figure 8:**
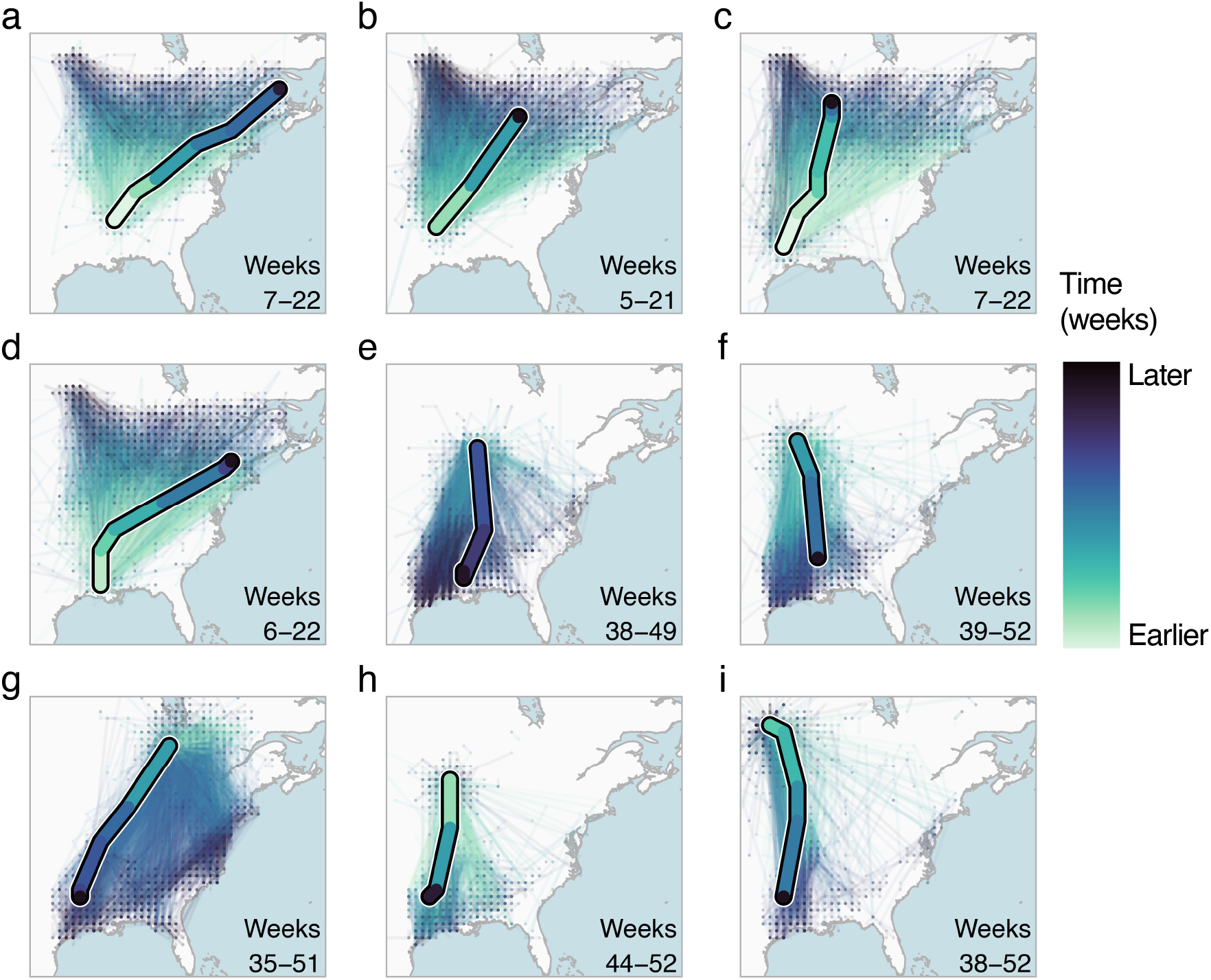
Model-simulated trajectories for GPS-tracked American Woodcocks *Scolopax minor*. Observed movements of GPS-tracked woodcocks (single thick path) and simulated trajectories (thin paths) for 2500 simulated birds originating at the same starting location as observed birds.

**Figure 9:**
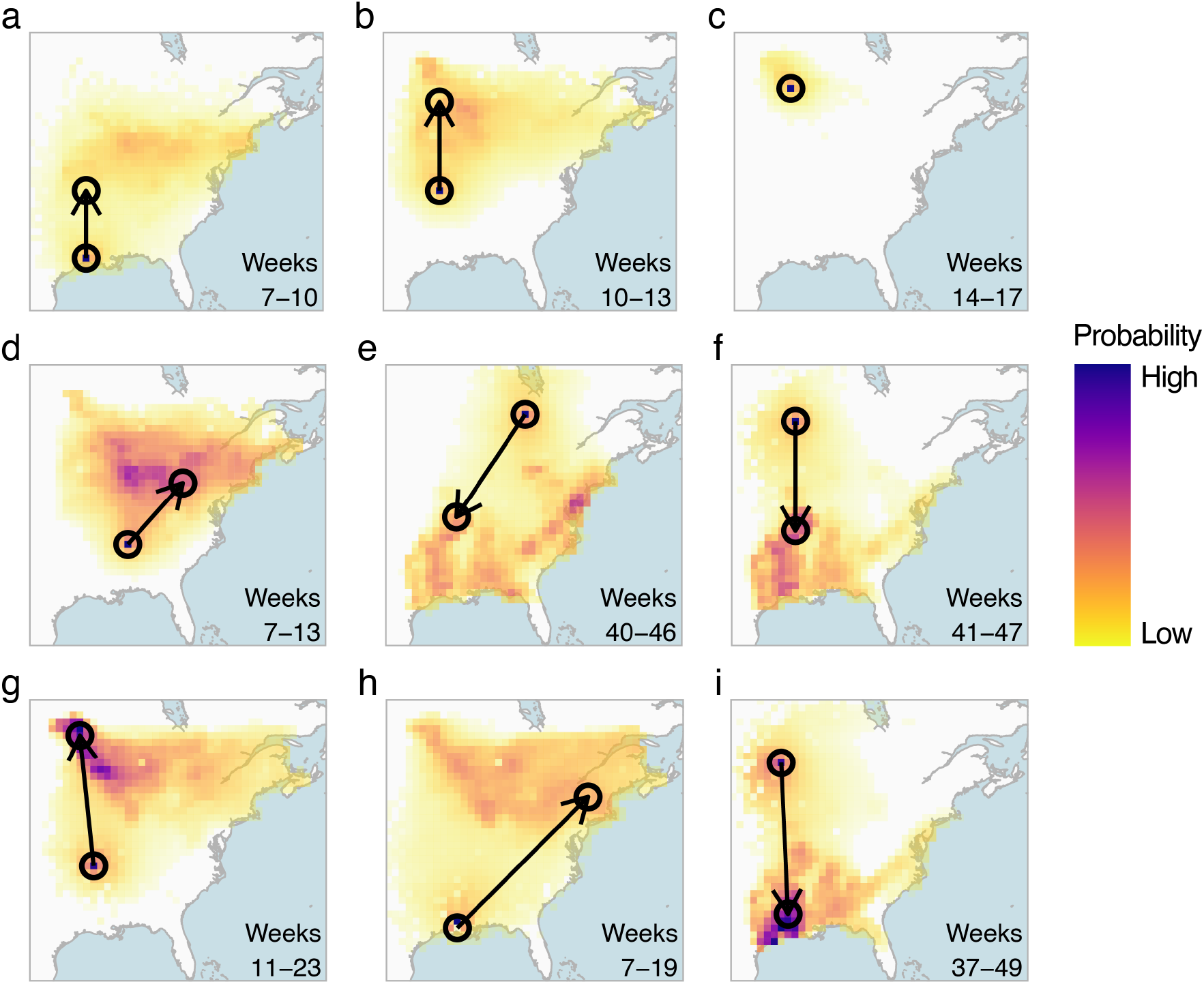
Conditional forecast distributions for GPS-tracked American Woodcocks *Scolopax minor*. Each heatmap shows the predicted movement distribution of a GPS-tracked individual originating within the circle at the base of the arrow. Darker colors indicate a higher predicted likelihood of movement to that area. The point of the arrow shows the observed ending location. Shown are examples of 3-week (a-c, same individual), 6-week (d-f, different individuals), and 12-week (g-i, different individuals) conditional forecasts.

## E Species Maps

**Figure 10:**
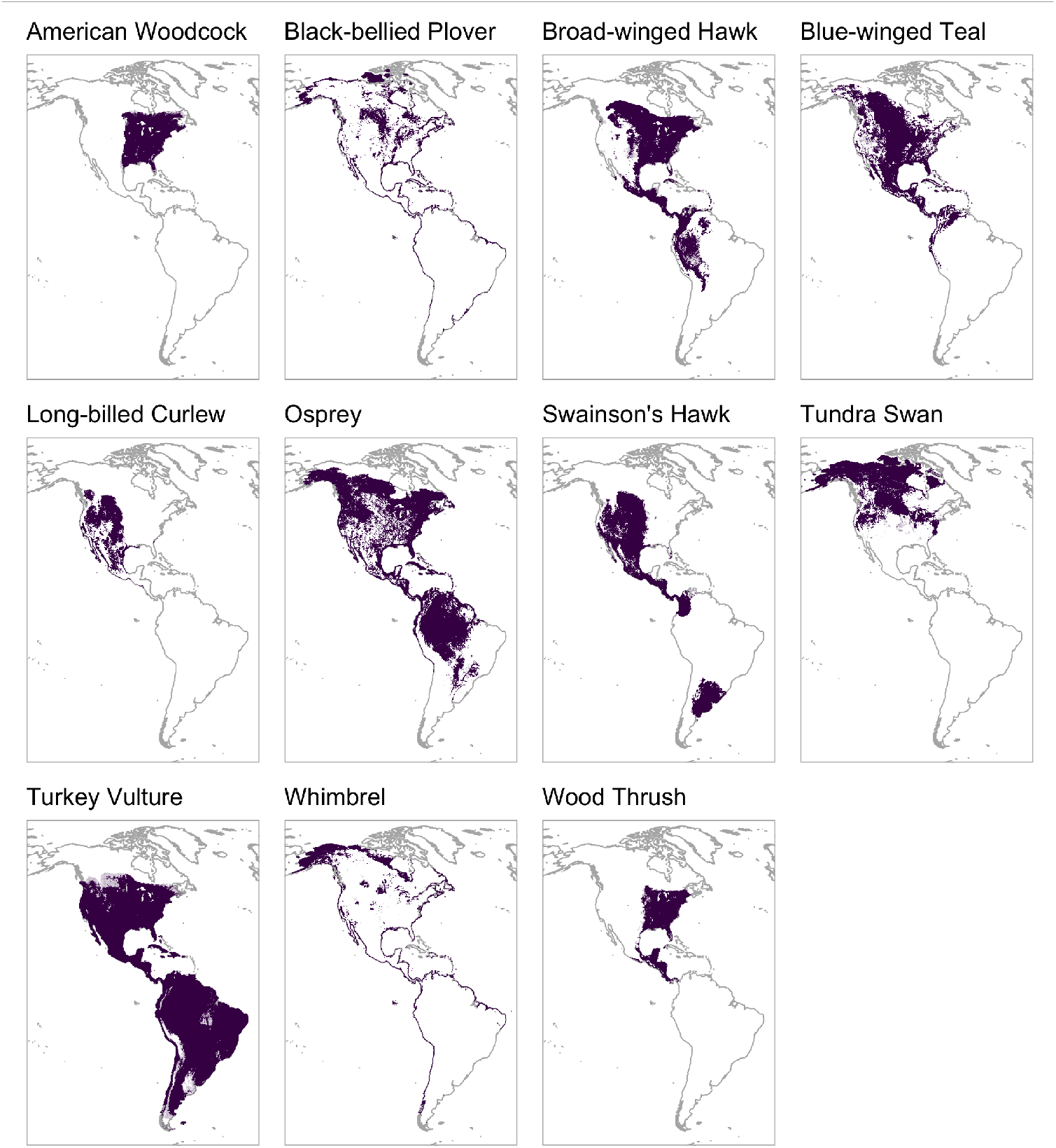
Geographic distributions of species modeled using eBird Status & Trends. To produce these plots, we standardized relative abundance outputs each week by dividing by the sum of total relative abundance, and then averaged these values over the entire year. Finally, we colored cells with the top 90% of values to show the species’ geographic distributions.

## Footnotes

1 https://ebird.org/science/status-and-trends

2 It is common in the literature to restrict to *time-homogeneous* Markov chains, where the transition probabilities are the same in every time step; we do not make this restriction.

